# Investigating the phylogenetic history of toxin tolerance in mushroom-feeding *Drosophila*

**DOI:** 10.1101/2023.08.03.551872

**Authors:** Theresa Erlenbach, Lauren Haynes, Olivia Fish, Jordan Beveridge, Eunice Bingolo, Sarah-Ashley Giambrone, Grace Kropelin, Stephanie Rudisill, Pablo Chialvo, Laura K. Reed, Kelly A. Dyer, Clare Scott Chialvo

## Abstract

Understanding how and when key novel adaptations evolved is a central goal of evolutionary biology. Within the *immigrans-tripunctata* radiation of *Drosophila*, many mushroom-feeding species are tolerant of host toxins, such as cyclopeptides, that are lethal to nearly all other eukaryotes. In this study, we used phylogenetic and functional approaches to investigate the evolution of cyclopeptide tolerance in the *immigrans-tripunctata* radiation of *Drosophila*. We first inferred the evolutionary relationships among 48 species in this radiation using 978 single copy orthologs. Our results resolved previous incongruities within species groups across the phylogeny. Second, we expanded on previous studies of toxin tolerance by assaying 16 of these species for tolerance to α-amanitin and found that six of these species could develop on diet with toxin. Third, we examined fly development on a diet containing a natural mix of toxins extracted from the Death Cap *Amanita phalloides* mushroom. Both tolerant and susceptible species developed on diet with this mix, though tolerant species survived at significantly higher concentrations. Finally, we asked how cyclopeptide tolerance might have evolved across the *immigrans-tripunctata* radiation and inferred that toxin tolerance was ancestral and subsequently lost multiple times. Our results suggest the evolutionary history of cyclopeptide tolerance is complex, and simply describing this trait as present or absent does not fully capture the occurrence or impact on this adaptive radiation. More broadly, the evolution of novelty can be more complex than previously thought, and that accurate descriptions of such novelties are critical in studies examining their evolution.

## Introduction

Evolutionary novelties involve sets of genetic and phenotypic changes that can permit a species to enter a new niche, provide release from selective pressures like competition, or initiate adaptive radiations (Simpson, 1953, Heard & Hauser, 1995). Most research on adaptations has focused on the evolution of structures, for instance novel body parts (Ohde et al., 2018, Hu et al., 2019, Broeckhoven et al., 2016). Less studied are biochemical adaptations, though these are likely as important for the processes of adaptation and speciation in many organisms. Some key biochemical adaptations are venom production, chemical sequestration, and toxin tolerance (Karageorgi et al., 2019, Giorgianni et al., 2020, Reimche et al., 2022).

A first step in understanding the emergence of a novel trait is to infer its phylogenetic history, meaning when did it evolve in the lineages where it is present. This inference requires a clear definition of what the trait is and a well-resolved phylogeny. Frequently, novel traits are treated as a binary state (*i.e.*, present/absent) in phylogenetic analysis, and this is typically clear for morphological traits (Tomita et al., 2021, Linz & Moczek, 2020). However, making this determination can be more difficult for biochemical traits where the threshold for determining presence/absence depends on the type of biochemical novelty being considered (Zaspel et al., 2014, Karageorgi et al., 2019, Kazandjian et al., 2021). For example, toxin tolerance is usually defined by whether taxa can successfully feed or develop on toxic hosts (Le Pelley, 1973, Jaenike et al., 1983). In many cases, lab-based assays are used as a proxy to determine toxin tolerance. However, the level of toxin tolerance may vary within and across species, thus a simple presence/absence model may underestimate the biological complexity of a biochemical trait.

Here we focus on the tolerance of certain *Drosophila* species to cyclopeptides, which is the deadliest class of mushroom toxin (Wieland 1986; Bresinsky and Besl 1990; Tang et al 2016). These toxins occur in some species of *Amanita* mushrooms, including the Death Cap and Destroying Angel, and are lethal to most eukaryotes, including humans. Fatalities are attributed to the amatoxin subclass, which act by binding to RNA polymerase II (RNAP II) and inhibiting mRNA production, leading to death (Lindell et al., 1970). Intriguingly, at least 12 *Drosophila* species in the *immigrans-tripunctata* radiation use mushrooms containing cyclopeptide toxins as developmental hosts (Scott Chialvo & Werner, 2018). A species is defined as ‘tolerant’ if they can survive from egg to adult eclosion on a laboratory diet containing 50 µg/g α-amanitin (Jaenike et al., 1983, Stump et al., 2011), though the toxic mushrooms these flies feed and develop on can contain up to 1600 µg of α-amanitin per gram of dried mushroom (Wieland, 1968). Studies have shown that while flies survive without visible impacts to fitness on the mean α-amanitin (250 µg/g) concentration, the extreme concentrations sometimes found in mushrooms (750 to 1000 µg/g) begin to affect the flies deleteriously (Jaenike, 1985).

The tolerance of mushroom-feeding *Drosophila* to cyclopeptides is different from many other insect biochemical adaptations. First, typically only specialist feeders use hosts containing highly toxic compounds (Whittaker & Feeny, 1971, Cornell & Hawkins, 2003). Mushroom-feeding *Drosophila*, however, are generalists that use a variety of fleshy non-toxic and toxic fungi, and in some cases also use fruit and vegetation as developmental hosts (Lacy, 1984, Jaenike & James, 1991). Second, insects feeding on toxic hosts are often reported to be insensitive to the toxic compounds due to target site mutations (Holzinger & Wink, 1996, Labeyrie & Dobler, 2004, Karageorgi et al., 2019), but in toxin tolerant *Drosophila*, the target of amatoxins, RNAP II, does not contain mutations that prevent binding (Jaenike et al., 1983, Stump et al., 2011). It was found though that inhibition of Cytochrome P450s resulted in decreased tolerance in four of eight species assayed (Stump et al., 2011), suggesting P450s may be important to the mechanism of tolerance in some species. Third, the gut microbiome can contribute to detoxifying secondary metabolites encountered by some insects (Ceja-Navarro et al., 2015, Shukla et al., 2018). In one species of α-amanitin tolerant flies, however, alteration of the larval gut microbiome did not result in loss of tolerance (Griffin & Reed, 2020). Thus, these common mechanisms do not appear to be responsible for toxin tolerance in mushroom-feeding *Drosophila*.

Here we study the evolution of cyclopeptide tolerance in mushroom-feeding *Drosophila* using phylogenetic and functional approaches. The evolutionary relationships in the *immigrans-tripunctata* radiation are poorly understood and vary depending on the species sampled and data used to construct the phylogeny (Spicer & Jaenike, 1996, Perlman et al., 2003, Dyer et al., 2011, Hatadani et al., 2009, Morales-Hojas & Vieira, 2012, Scott Chialvo et al., 2019, Finet et al., 2021). Therefore, we first inferred the evolutionary relationships among an increased sampling of 48 species in the *immigrans-tripunctata* radiation. We then expand upon previous studies that examined the effect of a single cyclopeptide toxin, α-amanitin (Jaenike et al., 1983, Jaenike, 1992, Stump et al., 2011) and assessed how consuming a diet containing a natural mix of toxins extracted from *A. phalloides* (Death Caps) impacts survival of both cyclopeptide tolerant and susceptible *Drosophila*. We combine these empirical data with our species tree to construct a hypothesis of the ancestral state of toxin tolerance across this radiation. In sum, our results suggest cyclopeptide tolerance arose once in the radiation and was then lost multiple times. Further, toxin tolerance in this radiation appears to be more complex than previously thought, with susceptible species being able to survive on diet with toxin.

## Methods

### Fly Stocks, Taxon Sampling and RNA Sequencing

We maintained fly stocks used in phylogenetic analyses and toxin tolerance assays at 22.5°C with 50% relative humidity and a 12:12 light:dark cycle. Flies were maintained on 4-24 Instant Drosophila Media (Carolina Biological) with the addition of a piece of mushroom (*Agaricus bisporus*) and a dental cotton roll.

The 48 species used in the phylogenetic analyses are listed in Table S1. The sampling includes all four *testacea* group species and 18 of the 26 *quinaria* group species. There are two strains of *D. subquinaria*, one inland (Hinton, Alberta) and one coastal (Portland, Oregon), as these populations are known to be strongly isolated (Jaenike et al., 2006). Members from the *immigrans*, *tripunctata*, and *cardini* groups were also included. The two outgroup species were *D. grimshawi* and *D. virilis.* The *D. grimshawi, D. pruinosa, D. virilis,* and *D. quadrilineata* genomes were obtained from the 101 Drosophilid Genomes Project (Kim et al., 2021). The *D. innubila* genome assembly was downloaded from NCBI (GenBank accession GCA_004354385.2). For the remainder of the species (Table S1) we generated transcriptomes. Flies were collected 24 hours after emergence, and RNA was extracted using the Omega Bio-Tek EZNA Total RNA Kit 1 using an equal number of flash frozen males and females. Samples were sent to NovoGene for library prep and sequencing. Library preparation included poly-A enrichment, and libraries were sequenced as 150 bp paired-end reads.

### Transcriptome Assembly and Dataset Generation

Transcriptomes were assembled using Oyster River Protocol v2.2.2 (MacManes, 2018) with default parameters. Each transcriptome and genome was assessed for completeness using the *metazoan* BUSCO v3.0.2 dataset v9 (Simao et al., 2015). We used TOAST (Wcisel et al., 2020) to generate an individual fasta file of each BUSCO ortholog, using the functions ParseBuscoResults and ExtractBuscoSeqs in R v3.6.1 (R Core Team 2019). For species with genomes (Table S1), we extracted the corresponding sequence for each ortholog from the genomes using samtools faidx v1.10 (Li et al., 2009). These sequences were appended to the ortholog fasta files generated from TOAST. The new fasta files for BUSCO orthologs were run through TOAST’s MafftOrientAlign, MissingDataTable, and SuperAlign functions to generate an alignment file for each of the 978 individual orthologs and a concatenated alignment file containing all orthologs (506,575,152 bp).

### Phylogenetic Analyses

We generated species trees using maximum likelihood and coalescent-based frameworks. To generate a maximum likelihood tree, we used the concatenated dataset generated above. We ran Model Finder (Kalyaanamoorthy et al., 2017) in IQ-Tree v2.0.6 (Nguyen et al., 2015) to determine the model that best supported the data and generated a maximum likelihood species tree using that best-fit model. We assessed branch support with 100 bootstrap replicates. To estimate the coalescent-based tree, we first generated a gene tree for each ortholog using RAxML v8.1.12 (Stamatakis, 2014) with the GTRCAT model and 100 bootstraps. Bootstrapped gene trees were then used to estimate the species tree in ASTRAL-III v5.6.1(Zhang et al., 2018). All trees were visualized using Figtree v1.4.4 (http://tree.bio.ed.ac.uk/software/figtree/).

Since the loci used and site rate variation can influence a tree topology, we performed a sensitivity analysis (Dowdy et al., 2020, Buddenhagen et al., 2016). Briefly, sites in each ortholog were placed into 10 bins based on evolutionary rate using TIGER v2.0 (Cummins & McInerney, 2011). The first bin contained invariant sites, the last bin the fastest-evolving sites, and the remaining sites were partitioned into the middle eight bins based on rate. The program AMAS (Borowiec, 2016) was used to concatenate the binned sequences for each ortholog. For each ortholog, we created eight alignments by sequentially adding more rapidly evolving bins (e.g. 2+3, 2+3+4, etc.); the first bin was excluded since it only included invariant sites. Using RAxML v8.1.12 (Stamatakis, 2014), maximum likelihood trees were generated for each alignment using the GTRCAT model and 100 bootstraps. We estimated the pairwise distance among trees using treeCMP v2.0.76 (Bogdanowicz et al., 2012). These trees were plotted using cmdscale in R v3.6.1 (R Core Team 2019), and the Euclidean distance of each subsampled tree to their average center was calculated. We removed loci that did not have sites in all of the original site bins. The remaining 489 loci were then ranked based on this distance and placed equally into eight inclusion sets, from 61 to 489 loci, with each successive set containing more outlier trees. This was performed for each of the eight binning subsets to create a total of 64 locus-inclusion sets. New alignments were generated for each locus-inclusion set, and these were used to create a maximum likelihood tree for each locus in RAxML v8.1.12 (Stamatakis, 2014) with bootstrap support values. These bootstrapped trees were used in ASTRAL-III v5.6.1 (Zhang et al., 2018) to identify support for the coalescent species tree for each locus-inclusion set. To assess support for the maximum likelihood tree, we concatenated the alignment for each inclusion set, and then used the RAxML -b flag to assess bootstrap support for each inclusion set. Heatmaps for each node of the tree were generated to show how support varies across these inclusion sets with the R package ggplot2.

### Toxin tolerance assays

We conducted three toxin tolerance experiments in which we reared larvae on diets with or without toxin and assessed survival to adulthood. All tolerance assays were conducted in 7.5mL glass scintillation vials containing 250 mg of a mixture consisting of 73.5% Instant *Drosophila* food and 26.5% ground freeze dried *Agaricus bisporus* mushroom resuspended in either 1mL of water or 1mL of toxin in water. A 1.5cm x 4cm piece of cotton watercolor paper was added as a pupation substrate. Each replicate consisted of placing early first instar larvae (15-25 per species; Table S2) into the vial and observing survival to adult for 30 days, and we conducted five replicate vials of each treatment.

Previous studies of *Drosophila* cyclopeptide tolerance (Jaenike et al., 1983, Spicer & Jaenike, 1996, Stump et al., 2011) characterized tolerance to ⍺-amanitin in 20 species distributed across six species groups in the *immigrans-tripunctata* radiation. To conduct a more comprehensive examination of the evolution of ⍺-amanitin, we assessed tolerance to ⍺-amanitin in 16 additional species from six species groups (Table S2). The larvae were reared on diets without toxin or with 50µg/g ⍺-amanitin (62.5µg ⍺-amanitin in 1mL water), which was the concentration used previously to identify tolerant species (Jaenike et al., 1983, Stump et al., 2011). We coded the survival of each individual larvae to adulthood using a binary strategy (0 = Dead; 1 = Survived). We quantified the impact of ⍺-amanitin on survival for each species separately using a generalized linear model with a binomial distribution, logit link, and bias-adjustment using the *brglm2* v0.8.2 package (Kosmidis, 2021). The only included model effect was toxin presence. Statistical analyses were completed in RStudio v2022.7.1.554.3 (RStudio Team 2022).

In the second two experiments, we assessed survival using a natural toxin mix (NTM). The natural toxins from *Amanita phalloides* mushrooms collected in December 2017 in Point Reyes, California were extracted as in Scott Chialvo et al (2020). The toxins were quantified using commercially available standards (e.g., ⍺-amanitin, β-amanitin, phalloidin, and phallacidin). We first used this mix to characterize the impact of amatoxin on survival by rearing larvae of three amatoxin tolerant species and eight susceptible species (Table S3). First instar larvae were placed on diets containing different concentrations of NTM in standard lab food, including no toxin, 5, 10, and 50μg/g amatoxins. As only amatoxins are known to pass through the gut lining of mammals (Köppel, 1993), we based concentrations used in these survival assays on the combined concentrations of ⍺-amanitin and β-amanitin in the extract (0.215µg/mL and 0.326µg/mL respectively). Thorax length of emerging adults was measured using ImageJ software (Schneider et al., 2012). Additionally, for five toxin species susceptible to ⍺-amanitin (either found by us or others to be susceptible), we performed additional assays to identify the maximum survivable amatoxin concentration (Table S3). Based on preliminary assays, four species were reared on six diets (0, 10, 15, 20, 25, and 30µg/g amatoxins) and the remaining species (*D. multispina*) was reared on five diets (0, 10, 20, 30, and 40µg/g amatoxins).

To quantify the effect of natural toxin mix on survival, we implemented a Bayesian generalized linear mixed model with a *Gaussian* standard deviation of 3 using the *blme* v1.0-5 package (Chung et al., 2013). We compared survival on diets containing different levels of natural toxin mix using a Tukey’s post-hoc test implemented in *multcomp* v1.4-20 (Hothorn et al., 2008). We compared the thorax length of flies that emerged from natural toxin mix diets containing 0, 5, 10, and 50µg/g amatoxins using a one-way ANOVA followed by a Tukey-Kramer HSD test (Table S3). For each species, we analyzed thorax length by sex; we excluded a sex if fewer than three adults of that sex emerged for a given treatment.

### Ancestral State Reconstruction of Toxin Tolerance

To analyze how toxin tolerance has evolved along the *immigrans*-*tripunctata* radiation, we inferred the ancestral state of toxin tolerance using RASP4 (Yu et al., 2020). This tool uses character state information for each species to infer the most likely ancestral state of each internal node of a phylogenetic tree. Data on toxin tolerance used for character state information is either from this study or previous studies (Jaenike et al., 1983, Spicer & Jaenike, 1996, Stump et al., 2011). Our categories were (A) at least 10% absolute survival on diet with 50 µg/g ⍺-amanitin or (B) less than 10% absolute survival on diet with 50µg/g ⍺-amanitin. We used the Bayesian Binary MCMC (BBM) analysis in RASP with a fixed Jukes-Cantor model and null root distribution for 5,000,000 generations using four chains sampled every 100 generations. The first 1,000 samples were discarded as burn-in. We used only species for which we had physiological data from ⍺-amanitin (there is too little sampling with natural toxin mix for a meaningful multi-state analysis) and pruned the other species from the phylogeny, resulting in a phylogeny with 35 species.

## Results

### Phylogenetic Inference

To resolve the species tree for the *immigrans-tripunctata* radiation, we generated transcriptomes or collected genomes for 48 species (Table S1). The final dataset contained 978 single-copy orthologs from the BUSCO *metazoan* database. If a locus was duplicated, we used the highest scoring sequence. For our transcriptomes, the average number of single-copy orthologs was 67.1%, with a duplicate average of 30.4%. The genomes had an average of 96.3% single-copy orthologs and 1.8% average for duplicates. The percentage of all BUSCO categories (Complete and single-copy, Complete and duplicated, Fragmented, and Missing) in each species are in Table S1.

We used maximum likelihood and coalescent methods to generate a species tree from the 978 orthologs. The best model for our maximum likelihood tree was GTR+F+R10. Both the maximum likelihood topology (Figure 1A) and coalescent analysis (Figure 1B) produced five main well-supported clades, which we refer to as clades A through E. Clade A consists of the *immigrans* group along with *D. pruinosa* and is sister to Clades B-E. Clade B is the *funebris* group. Clade C is the *quinaria* group and contains two sister clades within it (C1 and C2). Clade D is the *testacea* group and *D. bizonata*. Clade E represents the *cardini* and *guarani* groups, along with *D. tripunctata*, *D. macroptera*, and *D. pallidipennis*. Within each clade, species relationships are identical between the two topologies other than the relationship of *D. brachynephros* and *D. phalerata* within Clade C1. Among deeper nodes, the relationships of Clades B, C, D, and E differs between the two phylogenies, with nearly all these basal nodes well supported in both trees.

**Figure 1:**
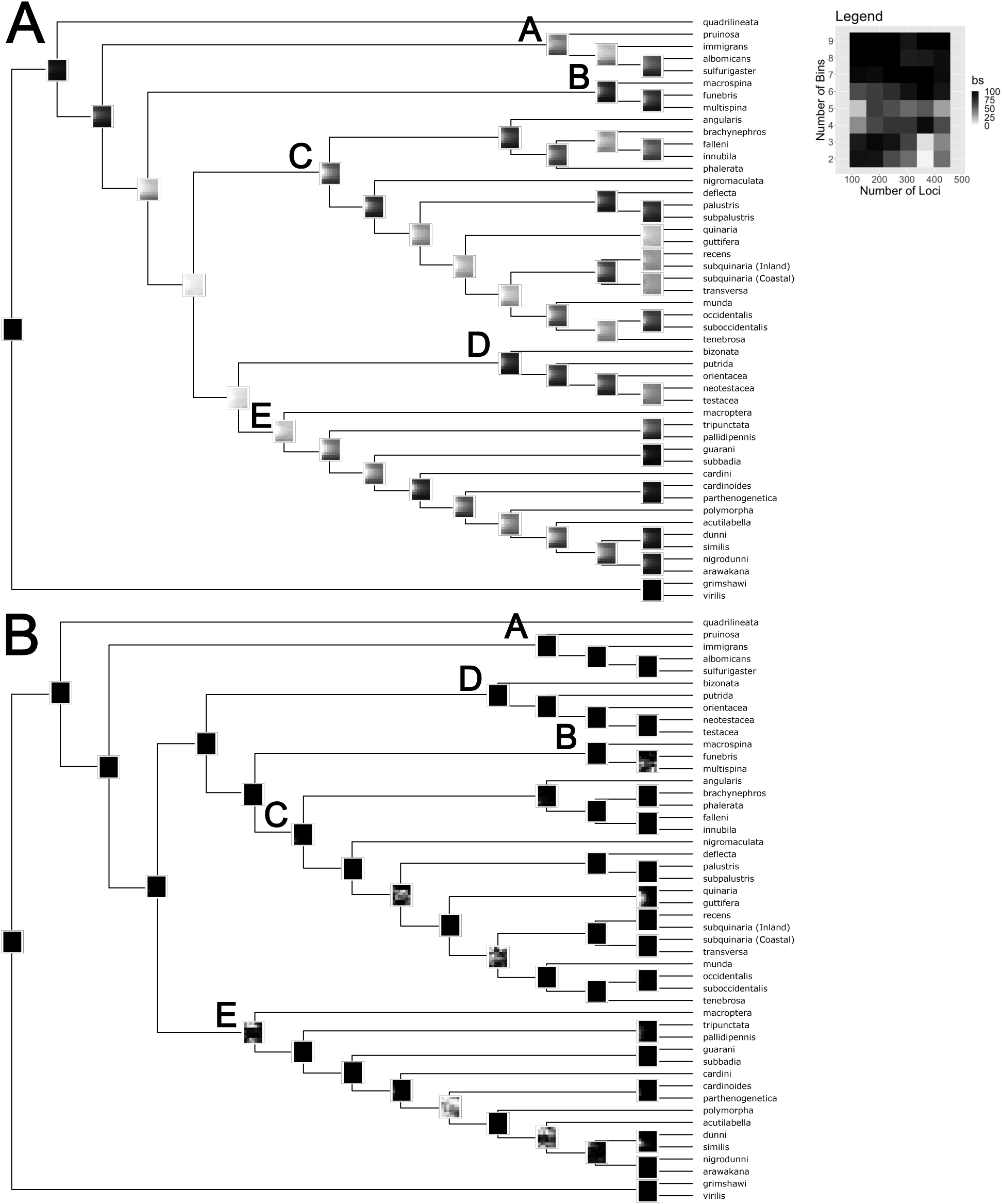
Species relationships for the *immigrans-tripunctata* radiation generated by (A) a maximum likelihood or (B) coalescent methods. Branch labels indicate bootstrap support and are only shown for branches with less than 100% support. Clade A in yellow represents the *immigrans* group and *D. pruinose,* Clade B in blue is the *funebris* group, Clade C in orange is the *quinaria* group, with two subclades marked C1 and C2, Clade D in purple is the *testacea* group and *D. bizonata*, and Clade E in green is the *cardini* and *guarani* groups along with *D. tripunctata*, *D. macroptera*, and *D. pallidipennis*.

We performed a sensitivity analysis to identify how the inclusion of faster-evolving loci affects support of our phylogenies and determine the best species tree for further analyses. The sensitivity analysis found substantial variability for support of internal nodes in our maximum likelihood tree (Figure S1A), with most nodes having much lower bootstrap support, especially those leading to clades C and D+E. In contrast, the coalescent phylogeny produced consistently higher bootstrap support values for most nodes independent of how many loci were included (Figure S1B). A few internal nodes had variability in bootstrap support, but the inclusion of more loci and variable sites increased their bootstrap support. This suggests that for tree construction using a coalescent approach, fewer loci are necessary to resolve a species tree, and maximum likelihood approaches require larger loci sampling with site rate variation to obtain the same level of resolution. Thus, we concluded the coalescent topology is the best supported species tree and use this phylogeny in remaining analyses.

### Toxin Tolerance

We assayed ⍺-amanitin tolerance in 16 species, representing six species groups, that had not previously been tested. We reared larvae on diets with and without ⍺-amanitin (50 µg/g) and measured the proportion that survived to adulthood (Figure 2, Table S2). In six species, at least 10% of larva survived to adulthood on the diet with toxin; these include *D. macrospina* in the *funebris* group, *D. guarani* and *D. subbadia* in the *guarani* group, and *D. occidentalis*, *D. suboccidentalis*, and *D. tenebrosa* in the *quinaria* group. Seven species produced no adults on ⍺-amanitin treatment, which significantly reduced their survival (*P*<0.0001). The remaining three species in the *cardini* group (*D. cardinoides, D. nigrodunni, D. parthenogenetica*) had greater than zero survival on ⍺-amanitin diet, but this was less than the 10% survival threshold.

**Figure 2:**
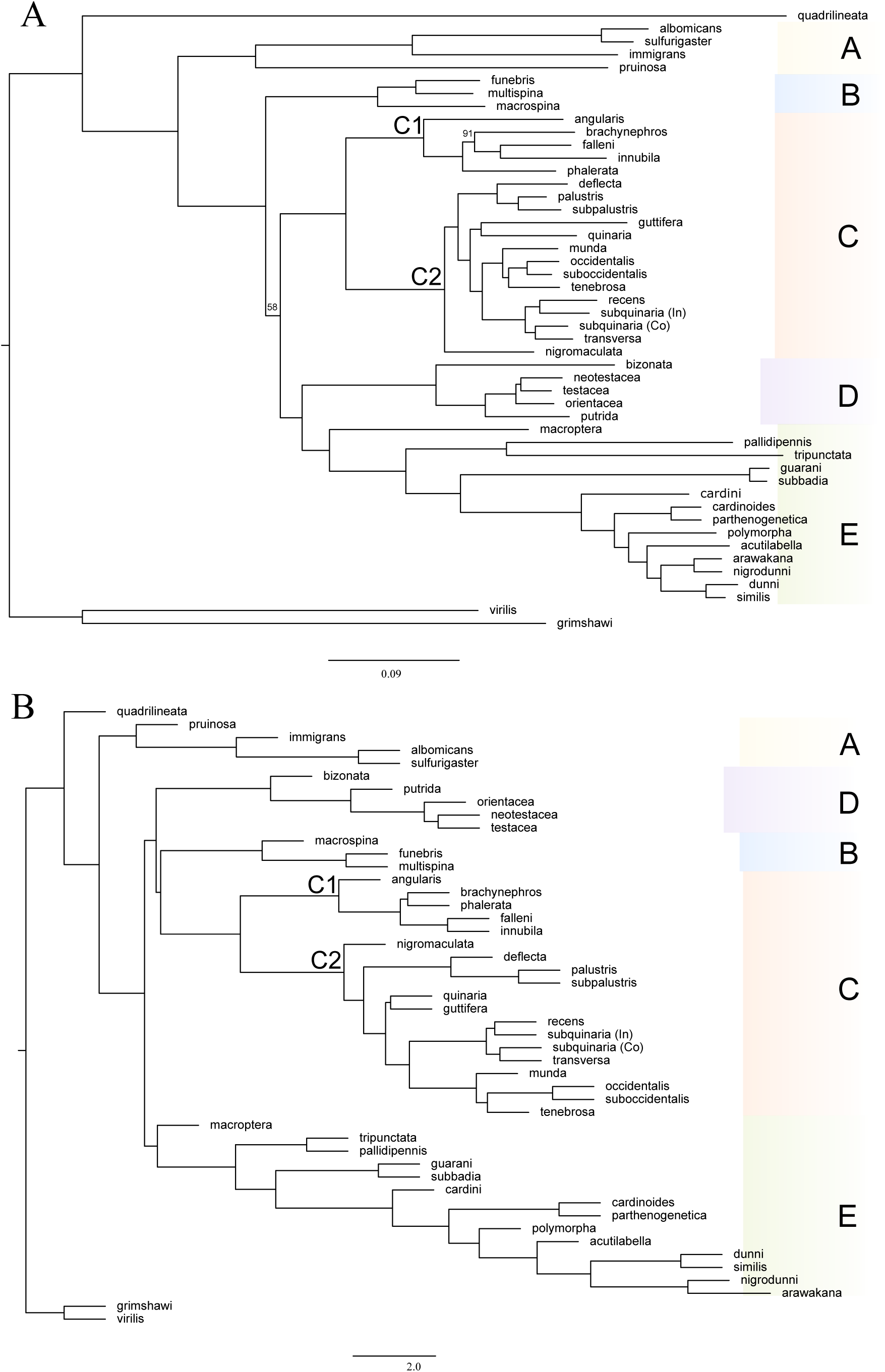
Proportion of first instar larvae that survived to adulthood on diets containing 0 or 50µg/g ⍺-amanitin. Species are clustered based on their species group and the error bars indicate the 95% binomial confidence intervals.

We next examined the responses of three toxin tolerant and eight susceptible species to diets containing a mix of toxins extracted from the toxic host *Amanita phalloides*. We measured both survival to adulthood (Figure 3; Table S3) and thorax length (Table S3). The three tolerant species represent three species groups (*quinaria*, *testacea*, and *tripunctata* groups); each of these species had high survival on all toxin concentrations included in the assay (Figure 3), and thorax length did not differ between treatments (Table S3). In *D. orientacea* (*testacea* group), survival was independent of toxin presence or concentration (*P* > 0.05). For *D. tripunctata*, larvae survived at higher numbers on each toxin diet, but the difference was only significant on the 5 and 10µg/g concentrations, while *D. recens* (quinaria group) had a significantly higher survival on diet containing the highest concentration of toxin (*P*=0.01).

**Figure 3.**
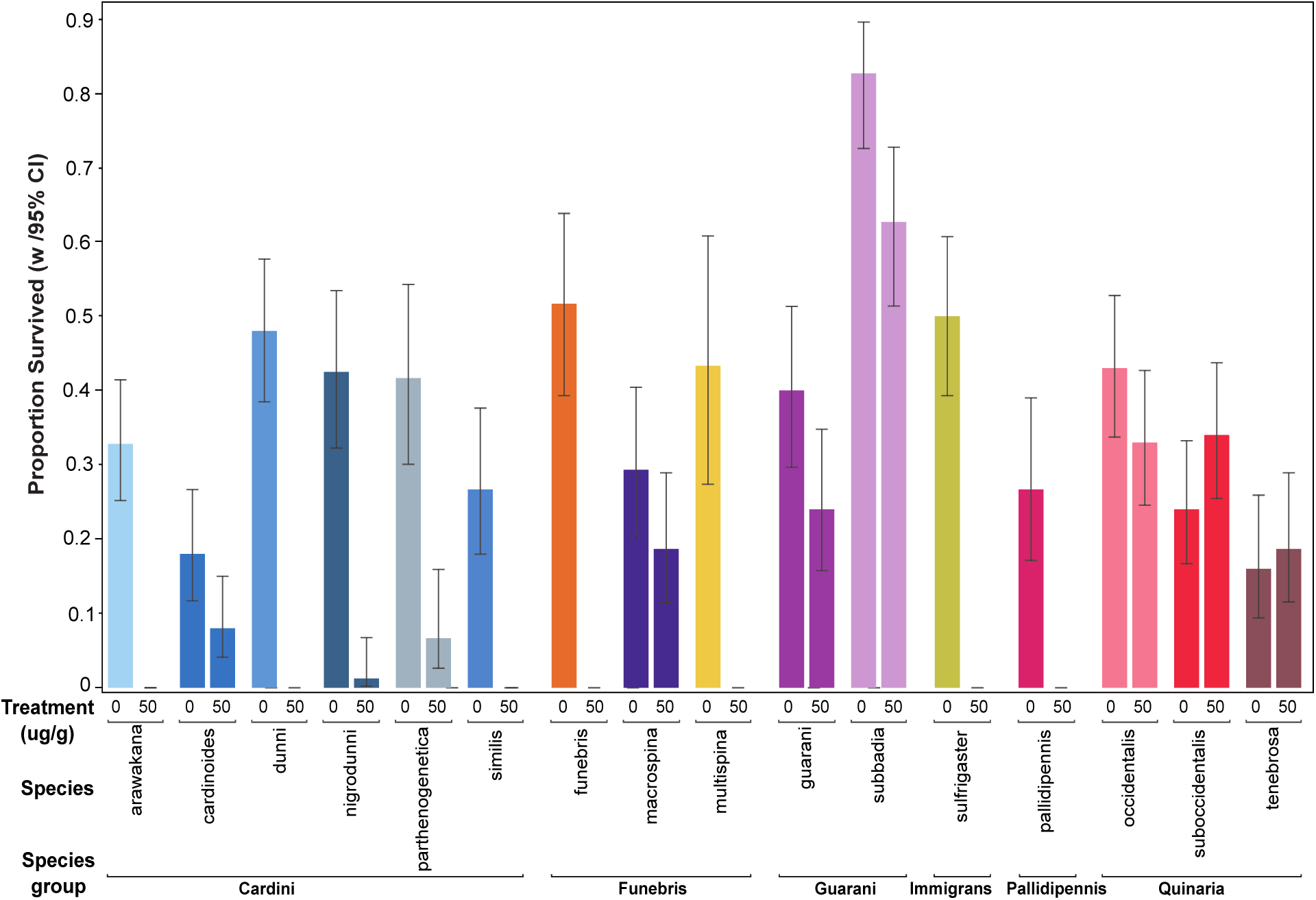
Proportion of first instar larvae that survived to adulthood on diets containing a range of concentrations of a natural mix of toxins extracted from *Amanita phalloides*. The error bars indicate the 95% binomial confidence interval. Mushroom image indicates that a species is known to be tolerant whereas skull and crossbones image denotes a species as being susceptible to toxin, based on previous work. *D. orientacea* is putatively tolerant because it breeds on mushrooms, but it has not been tested for tolerance to ⍺-amanitin. Species relationships are indicated by the phylogeny along the X-axis.

The susceptible species reared on diets with and without toxin mix included a single genotype of *D. melanogaster* (NC-1) and seven species from the *immigrans-tripunctata* radiation representing five species groups. *D. melanogaster* only survived on the lowest toxin concentration (5µg/g; *P* < 0.001; Figure 3), and females that survived were larger than controls (*P* < 0.05). The seven susceptible species from the *immigrans-tripunctata* radiation all survived on diets containing 5 and 10µg/g amatoxins (Figure 3). Relative to rearing on a diet without toxin, at these concentrations, four species did not have reduced survival (*P* > 0.05), *D. multispina* had higher survival (*P* < 0.001), and *D. funebris* and *D. pallidipennis* had reduced survival (*P* < 0.001 and P < 0.01, respectively). Interestingly, of these species only *D. funebris* had survival >5% on diet with 50µg/g amatoxins, though we note in our previous assay with pure ⍺-amanitin it did not survive at this concentration. The toxin diet only effected adult thorax length for *D. funebris* males, where individuals emerging on the two higher toxin concentrations (10 and 50µg/g) had longer thoraxes (P < 0.05) than the 5µg/g diet (Table S3).

We conducted additional experiments to narrow the range of the lethal dose of amatoxins in five of these susceptible species. We reared larvae on diet containing between 15 and 40µg/g natural amatoxins and scored survival and found that each species responded in a unique manner (Figure 3, Table S3). More than 10% of larvae of both *D. deflecta* (*quinaria* group) and *D. multispina (funebris* group) survived on diet with amatoxin up to 30µg/g. More than 10% of *D. pallidipennis* (*pallidipennis* group) survived on diet with amatoxin up to 15µg/g, but no adults emerged on diets with higher concentrations. In comparison, *D. dunni* (*cardini* group) and *D. immigrans* (*immigrans* group) showed decreasing survival with increasing toxin concentration.

### Evolution of Toxin Tolerance

With the knowledge that toxin tolerance varies within the *immigrans-tripunctata* radiation, we sought to reconstruct the ancestral state of this trait. We examined toxin tolerance as a binary trait, using results from our toxin tolerance assays and previous studies (Jaenike et al., 1983, Spicer & Jaenike, 1996, Stump et al., 2011). We define tolerant as at least 10% survival at 50 µg/g of ⍺-amanitin and susceptible less than 10% survival at this concentration. There is uncertainty on toxin ancestry for our deepest node in the tree, but nodes in the phylogeny leading to Clades B-E all indicate tolerance, which is subsequently lost at least five times (Figure 4).

**Figure 4:**
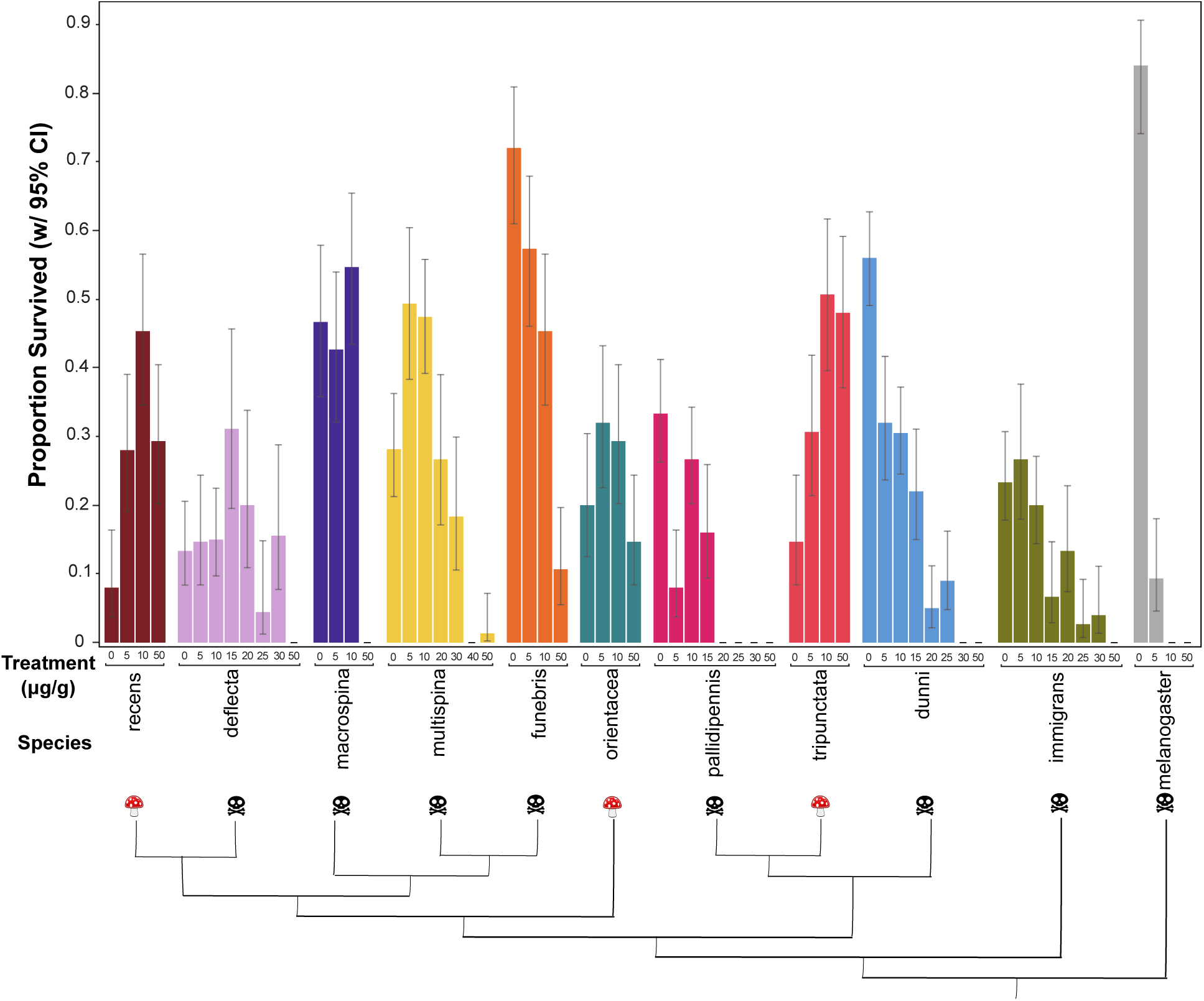
Ancestral state reconstruction of toxin tolerance evolution. The ancestral state of toxin tolerance is indicated on each branch by the circles, with the probability indicated by the colors, as shown in the legend. We also denote losses of tolerance along the tree.

There is one loss of tolerance in the *funebris* group (Clade B) and two in the *quinaria* group (Clade C), including one in *D. quinaria* and another the common ancestor of *D. deflecta*, *D. palustris*, and *D. subpalustris*. Toxin tolerance is more variable in Clade E, where about half of the species assayed are susceptible. Tolerance was lost on the branch leading to *D. pallidipennis* and, for the *cardini* group, was either lost twice or lost once and then regained, and we suggest the former is most parsimonious.

## Discussion

Species within the *immigrans-tripunctata* radiation exhibit a wide range of trait variation, including morphological characters, host preference, parasite prevalence, and toxin tolerance (Dombeck & Jaenike, 2004, Simunovic & Jaenike, 2006, Werner et al., 2010, Markow & O’Grady, 2008, Spicer & Jaenike, 1996, Stump et al., 2011). Of particular fascination is the tolerance found in many of these species to cyclopeptide toxins, as these are among the only eukaryotes known to consume and develop on these potent toxins (Scott Chialvo & Werner, 2018). Species that are generalist feeders on fleshy mushrooms or a combination of mushrooms and rotting fruits and/or vegetative matter are able to use cyclopeptide-containing mushrooms as developmental hosts (Jaenike et al., 1983, Lacy, 1984), while other more specialized mushroom feeders in the radiation (*i.e.*, *D. funebris*) are not (Stump et al., 2011, Korneyev, 2010). To broaden our understanding of cyclopeptide toxin tolerance and the evolutionary relationships within the *immigrans-tripunctata* radiation, we generated a transcriptome phylogeny, conducted a series of survival assays, and reconstructed toxin tolerance in the radiation.

When attempting to understand how traits among a lineage evolve, it is imperative to ensure that sufficient loci and a large enough taxon sampling are used in tree-building. Recently, research on the superfamily Ephydroidea, which contains Drosophilidae, sought to tease apart inconsistencies with a broader sampling of species and increased number of nuclear genes (Winkler et al., 2022). This work reaffirmed and supported the need for broad taxonomic sampling across a lineage to best understand the evolution of species and the traits that encompass them. Phylogenetic studies of the *immigrans-tripunctata* radiation have used either limited species sampling and/or few loci (Hatadani et al., 2009, Dyer et al., 2011, Izumitani et al., 2016, Scott Chialvo et al., 2019), resulting in incongruence among the recovered tree topologies. We reconstructed a species tree for 48 species in the *immigrans-tripunctata* radiation using maximum likelihood and coalescent phylogenetic approaches with 978 single-copy orthologs. Both trees produced the same five major clades, though there was variation in where they fell along the topology (Figure 1). Previous work using a smaller subset of this radiation could not confirm the monophyly of the *quinaria* group (Scott Chialvo et al., 2019). Our results support monophyly of the *quinaria* group, as the trees recovered from both analyses had the two subclades sister to one another (Figure 1). Interestingly, the *testacea* group (Clade D) has different locations along our topologies, either sister to Clade E (Figure 1A) or to the combined clades of B and C (Figure 1B). Due to these inconsistencies, we performed a sensitivity analysis and found the coalescent phylogeny produced the greatest support across all bin and loci combinations (Figure S1). With factors such as incomplete lineage sorting giving high support to an incorrect topology generated with a concatenated loci dataset (Kubatko & Degnan, 2007, Mendes & Hahn, 2018, Roch & Steel, 2015), it is not surprising the coalescent topology (Figure 1B) was the most consistent. Other published phylogenies show similar topologies, including the *testacea* and *funebris* groups sharing immediate ancestry (Finet et al., 2021, Zhang et al., 2021).

We surveyed 16 species from six species groups, six of which can survive at greater than 10% on a diet containing 50μg/g α-amanitin (Figure 2). Three are members of the *quinaria* species group, which includes many tolerant species (Jaenike et al., 1983, Spicer & Jaenike, 1996, Stump et al., 2011). The other three are from the *guarani* and *funebris* species groups. The *guarani* group is neotropical (Penafiel-Vinueza & Rafael, 2018), and very little is known about their natural history. In the *funebris* group, one of the three species tested, *D. macrospina*, survived at a high level on the ⍺-amanitin diet. This species breeds in woodlands near streams (Mainland, 1942, Miller et al., 2017), but nothing is known about its host usage. Another susceptible member of the group, *D. funebris*, is a specialized feeder on polypore, shelf fungi (Korneyev, 2010); this species did not produce adults on the 50ug/g α-amanitin diet.

Interestingly, we find variable tolerance in the *cardini* group. For example, there is weak survival (<10%) on 50μg/g α-amanitin diet for *D. cardinoides*, *D. parthenogenetica,* and *D. nigrodunni. Drosophila acutilabella*, another member of the *cardini* group, is known to feed on mushrooms and has higher survival on toxin food (21%)(Stump et al., 2011). Given that all previously tested α-amanitin tolerant species use fleshy mushrooms as hosts (Jaenike et al., 1983, Lacy, 1984), our results suggest cyclopeptide tolerance is associated with this specific feeding behavior, and the presence of toxin tolerance in a species could serve as an indicator of its host usage.

Including our results, cyclopeptide tolerance has been identified in seven species groups in the *immigrans-tripunctata* radiation based on whether larvae can survive on a diet containing 50µg/g ⍺-amanitin. While the 50µg/g concentration is below the mean ⍺-amanitin concentration (250µg/g) in *Amanita* mushrooms (Tyler Jr. et al., 1966, Yocum & Simons, 1977), only species that consume fleshy mushrooms can survive on this concentration (Jaenike et al., 1983, Stump et al., 2011). Thus, cyclopeptide tolerance is more broadly distributed within the *immigrans-tripunctata* radiation than previously known. Our ancestral state reconstruction of toxin tolerance supports this, where we find that toxin tolerance evolved early in the radiation and was lost multiple times (Figure 4). These results would also remain if we were to use the maximum-likelihood tree for reconstruction (data not shown). This is consistent with Spicer and Jaenike (1996), who proposed that toxin tolerance evolved once and was lost multiple times in the *quinaria* group. We note that Stump et al (2011) showed that inhibition of Cytochrome P450s affected toxin tolerant species differently but did support tolerance as the ancestral state.

Although toxic *Amanita* mushrooms can contain over 10 different cyclopeptide toxins, previous studies of toxin tolerance focused only on ⍺-amanitin (Jaenike et al., 1983, Jaenike, 1985, Jaenike, 1992, Stump et al., 2011). To fully understand a complex trait such as toxin tolerance, it is best to expose organisms to a diet that most closely resembles what they would encounter in the wild (Scott Chialvo et al., 2020). Therefore, we reared three tolerant and five susceptible species on diets containing a natural mix of toxins extracted from the Death Cap *Amanita phalloides* (Figure 3). As expected, the three toxin tolerant species survived at a high level on all toxin concentrations. Somewhat surprisingly, we found every toxin susceptible species we included from the *immigrans-tripunctata* radiation survived on 10µg/g amatoxins. In additional assays, the maximum survivable concentration differed among species and ranged from 15µg/g (*D. pallidipennis*) to 30µg/g (*D. deflecta, D. immigrans,* and *D. dunni*) amatoxins. Of the susceptible species, only *D. funebris* survived at 50µg/g amatoxins. Given this species did not survive on 50ug/g α-amanitin (Figure 2), this suggests there may be antagonistic interactions present within the toxin mix that impact toxicity. In addition, intraspecific variation in tolerance to the natural toxin mix is present in four mushroom-feeding *Drosophila* species (Kokate et al., 2022), including *D. neotestacea* (*testacea* group), which has not been tested for tolerance to ⍺-amanitin. *D. putrida* is tolerant of 50ug/g α-amanitin (Jaenike et al., 1983), and we showed that *D. orientacea* survives on the natural toxin mix, suggesting that the *testacea* group is tolerant of amatoxins.

In conclusion, our results suggest that evolution of amatoxin tolerance in the *immigrans-tripunctata* radiation is more complex than initially appreciated. Species display different levels of tolerance, and even species previously described as susceptible show some tolerance.

*Drosophila melanogaster* only survived at a low level at the lowest concentration of the natural toxin mix (Figure 3), suggesting this trait is not universal to *Drosophila.* The toxin concentration the susceptible species survived at in this experiment is likely higher than what these organisms experience in their natural environment, thus it is unclear what maintains this trait. These results imply toxin tolerance may represent a synapomorphy of the radiation, and the concentration of tolerance in species not regularly exposed to the toxin has reduced over time. In sum, the combination of our phylogenetic and functional approaches allowed us to analyze a complex biochemical trait and suggest a complete understanding of a trait is necessary to elucidate its evolution across an adaptive radiation. With regards to studies of the evolution of biochemical novelties, we suggest that classifying adaptations as present/absent and conducting assays with only one toxin may not accurately capture the full range of trait variation; thus, leading to oversimplified or inaccurate understandings of evolutionarily important traits.

## Acknowledgements

This work was funded by grants from the National Science Foundation (DEB173824 to KAD, DEB-1737869 to CSC and LKR, DBI-2217912 to CSC) and the National Institutes of Health (under T32GM007103 to TE).

## Supplemental Tables (Included as supplemental files)

**Table S1.** Species used in the study and their respective species groups. ⍺-amanitin tolerance is defined as (A) survival at 50 µg /g ⍺-amanitin or (B) no survival at 50 µg /g ⍺-amanitin. Natural toxin mix (NTM) tolerance is defined as survival at (A) 50 µg/g NTM, (B)≤ 30 µg/g NTM, (C) ≤25 µg/g NTM, (D) ≤20 µg/g NTM, (E) ≤15 µg/g NTM, and (F) ≤10 µg/g NTM. Feeding behavior is described as M: mushrooms, V: decaying vegetation, and F: fermenting fruit. Number and total length of reads are included for species sequenced in this study, as well as the type of assembly for all samples. BUSCO v9 scores are based on *metazoan* dataset. NT indicates ’not tested’ as we do not have data for these.

**Table S2.** Impact of diets containing α-amanitin on larvae to adult survival.

**Table S3.** Impact of diets containing natural toxin mix on larvae to adult survival to adult and thorax length. This table also indicates the maximum survivable concentration in susceptible species.

**Table S4.** Impact of toxin diet on thorax size.

**Figure S1:**
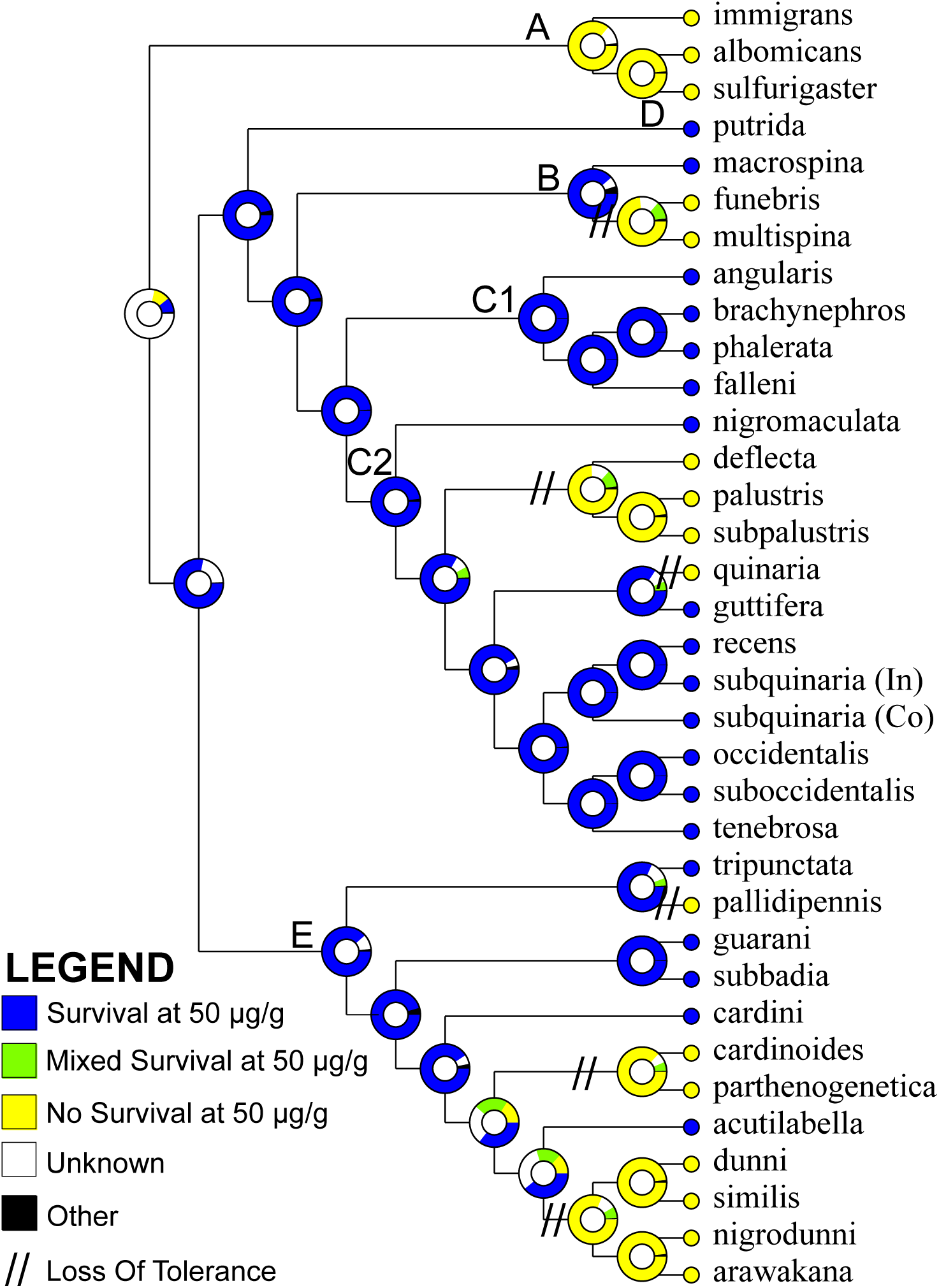
Sensitivity analysis for the (A) maximum likelihood species tree and (B) coalescent analysis species tree. Legend inset shows a randomly chosen heatmap, which gives the number of loci (X-axis) and number of bins (Y-axis) used. The color gradient indicates bootstrap support for the branches based on the loci and bin combinations in the sensitivity analysis, with white being 0 bootstrap support (bs) and black 100 bootstrap support. In (A), we find lower bootstrap support for intermediate bin combinations (4-6) at all sets of loci for many nodes. This can most likely be attributed to a large portion of the sites in the locus-inclusion set being largely conserved for bins 1-3, but the addition of more variable sites for intermediate bins (4-6), impacts tree reconstruction during bootstrapping. When most sites are conserved in lower bins (1-3), bootstrapping is going to produce similar topologies each time; however, when some of the sites in the alignment become variable as they are in intermediate bins, the resampling done during bootstrapping could produce topologies that vary from the main tree because there is random variation at a portion of the sites. Bootstrap support is recovered when we include more sites (bins 7-9), likely due to some of the variable sites being shared between closely related species and the resampling during bootstrapping occurring with a larger number of sites that are a more equal mix between conserved and variable.

## Notes

### Competing Interest Statement

The authors have declared no competing interest.

